# Cross-subgenus hybridization between *Leishmania* and *Sauroleishmania* informs on parasite genomic compatibility and transcriptomic adaptation

**DOI:** 10.1101/2025.03.25.645178

**Authors:** Viviane Noll Louzada-Flores, Pascale Pescher, Thomas Cokelaer, Tiago Rodrigues Ferreira, Maria Stefania Latrofa, Jairo Alfonso Mendoza-Roldan, Domenico Otranto, Gerald F Späth, Isabelle Louradour

**Affiliations:** Department of Veterinary Medicine, University of Bari, Bari, Italy; Institut Pasteur, Université Paris Cité, INSERM U1201, Unité de Parasitologie moléculaire et Signalisation, F-75015 Paris, France; Laboratory of Parasitic Diseases, National Institute of Allergy and Infectious Diseases, National Institutes of Health, Bethesda, MD, 20892, USA; Department of Veterinary Clinical Sciences, City University of Hong Kong, Hong Kong, China

**Keywords:** *Leishmania* hybrids, Sexual reproduction, *Leishmania infantum*, *Leishmania tarentolae*, Genome, Transcriptome

## Abstract

*Leishmania* parasites can enter a cryptic sexual reproductive cycle generating hybrid genotypes that can lead to unpredictable clinical outcomes and transmission cycles. Despite the importance of hybrids in *Leishmania* epidemiology, the mechanisms involved in their formation - including the impact of parental genetic distance - remain poorly understood. We report the *in vitro* generation of a hybrid between *Leishmania (Leishmania) infantum* and *Leishmania (Sauroleishmania) tarentolae,* two species from sister phylogenetic clades circulating across Southern Italy, providing evidence of genomic compatibility. Whole-Genome Sequencing indicates that, while the hybrid is largely tetraploid, its genome is not just the sum of its parental content. RNA-seq analysis of the hybrid transcriptome uncovers significant differences in the abundance of orthologous transcripts expressed from both parental genomes, driven by either parent-specific gene copy number variations or differential mRNA turnover. These results demonstrate that, beyond genomic restructuring, post-transcriptional regulation may serve as an additional mechanism shaping viable hybrid phenotypes, potentially enhancing parasite adaptability and fitness.

## Introduction

Unicellular eukaryotes can reproduce asexually through clonal reproduction, giving rise to a progeny that is genetically identical to the parental organism, and/or sexually, where two parental genomes are combined to create a new one. These forms of reproduction are not always exclusive, as for example observed in protists and fungi **[1–4]**. This applies to parasites of the genus *Leishmania,* the causative agents of leishmaniases in many animal species, including humans. These vector-borne diseases are worldwide distributed in tropical and sub-tropical regions and present a range of clinical manifestations, from self-limiting cutaneous lesions to visceral complications, the latter leading to death if untreated **[5]**. At least 20 different *Leishmania* species have been identified, infecting a large number of sand fly vectors and animal reservoirs, and showing different tissue tropisms and clinical outcomes **[6]**.

*Leishmania* parasites have a diphasic life cycle, with extracellular promastigotes multiplying asexually within the digestive tract of their insect hosts, the phlebotomine sand flies, and intracellular amastigotes that multiply asexually within the macrophages of the vertebrate hosts **[7]**. Clonal reproduction was for a long time regarded as the only reproductive strategy of *Leishmania* parasites. However, it is now clear that they can also enter a cryptic, sexual reproductive cycle leading to the production of hybrids, as first hypothesized by the observation of hybrid field isolates **[8–14]** and later demonstrated through experimental hybridization studies **[15]**. The possibility of selfing – a form of sexual reproduction where mating occurs between genetically identical parasites derived from the same individual – has recently been demonstrated in *Leishmania* spp., further challenging the notion of its clonal propagation **[16]**.

Facultative sex creates new genotypes with unpredictable phenotypes, which may lead to changes in tissue tropism, virulence, or the emergence of drug resistance **[2]**. This is the case of *L. infantum/L. donovani* hybrids, recently identified as the cause of re-emerging human leishmaniasis in the North of Italy **[17]**. Different strains and/or species of *Leishmania* frequently circulate in the same area, and co-infections have been recorded in different hosts, creating the possibility of hybrid generation during sand fly co-infection **[18]**. Both the description of mixed infections and natural hybridization represent a potential challenge to medical and veterinary practitioners, as they increase the possibility of diagnostic inaccuracies due to cross-reactivity in the detection tests **[19,20]**.

The saurian-associated *L. tarentolae,* infecting reptiles and transmitted through the bite of *Sergentomyia minuta* sand flies **[21,22]** is non-pathogenic to mammals. In contrast, *L. infantum* is transmitted during the blood meal of many sand flies in the genus *Phlebotomus* spp. **[23]** and is pathogenic to humans, where it can lead to visceral leishmaniasis **[7]**. Dogs are the main reservoirs of *L. infantum* and the infection may cause a range of clinical signs, including skin lesions, weight loss, lethargy, and swollen lymph nodes **[24]**. There are several lines of evidence that both *L. tarentolae* and *L. infantum* parasites may co-occur in the sand flies and possibly engage in hybridization. First, the recent molecular detection of *L. tarentolae* in human blood in central Italy **[25]** and by PCR and immunofluorescence antibody test (IFAT) in sheltered dogs in Italy **[19]** suggests that this parasite species could also circulate in mammals. Of note, no live *L. tarentolae* parasites were recovered from those dogs **[19,20]**. Second, human and dog blood was detected in engorged *S. minuta* females, showing that these flies do not feed exclusively on reptiles **[26]**. Third, *L. infantum* amastigote-like forms and its DNA was detected in lizards and geckoes residing in or close to dog shelters in southern Italy **[21]**. Finally, experimental infections demonstrated that *L. tarentolae* is also able to complete its life cycle in *P. perniciosus* **[27]**. While these arguments suggest the possibility for both species to be present simultaneously inside the same sand fly thus opening the possibility for hybrid formation **[19]**, no such extreme, inter-species hybrid has been so far identified in nature. This raises the question if such hybrids just escaped detection due to low frequency and limited sampling, or whether the genetic distance between the possible parent precludes the formation of viable hybrids.

In this study, we addressed this important open question and tested whether *L. tarentolae* and *L. infantum* were able to hybridize using an *in vitro* hybridization protocol **[28, 29]**. We present phenotypic, genetic, and transcriptomic evidence for successful production of a viable hybrid strain, thus providing first evidence for genetic compatibility between these two very different parasite species. Our results call for improved diagnostic tests that can identify *Leishmania* spp. hybrids in patient material and open new avenues of investigation regarding their mechanisms of adaptation and evolution.

## Results

### Generation of inter-subgenus *L. infantum*/*L. tarentolae* hybrids *in vitro*

To test whether *L. infantum* and *L. tarentolae* are able to hybridize, we set up *in vitro* crosses between a GFP+, neomycin-resistant *L. infantum* strain (Linf-GFP) and either an RFP+, hygromycin-resistant *L. tarentolae* strain (Ltar-RFP) or an RFP+, hygromycin-resistant *L. infantum* strain (Linf-RFP), used as a control for *in vitro* hybridization (**Figure 1A**). We performed 6 independent series of Linf/Ltar, and Linf/Linf crosses, representing a total of 504 culture wells for each of the two parent combinations (**Figure 1B**). Each cross series produced hybrids between Linf-GFP and Linf-RFP (a total of 9 hybrids, referred to as Linf/Linf), but only 1 of the 6 cross series gave rise to a single hybrid between Linf-GFP and Ltar-RFP (referred to as Linf/Ltar), reflecting a minimum frequency of mating-competent cells in the parental cultures ranging from 3.17E-9 to 1.32E-9 for the Linf x Linf and 2.16E-9 to 1.49E-9 for the Linf x Ltar crosses (**Figure 1B**). Flow cytometry confirmed the hybrid nature of the double-drug resistant parasites, which all simultaneously express the fluorescent markers of both parental strains (**Figure 1C**). These data provide first evidence of experimental F1 hybrid production between *L. infantum* and *L. tarentolae*, which seems a very rare event even *in vitro*, likely due to the strong genomic shock produced when hybridizing such distant species.

**Figure 1:**
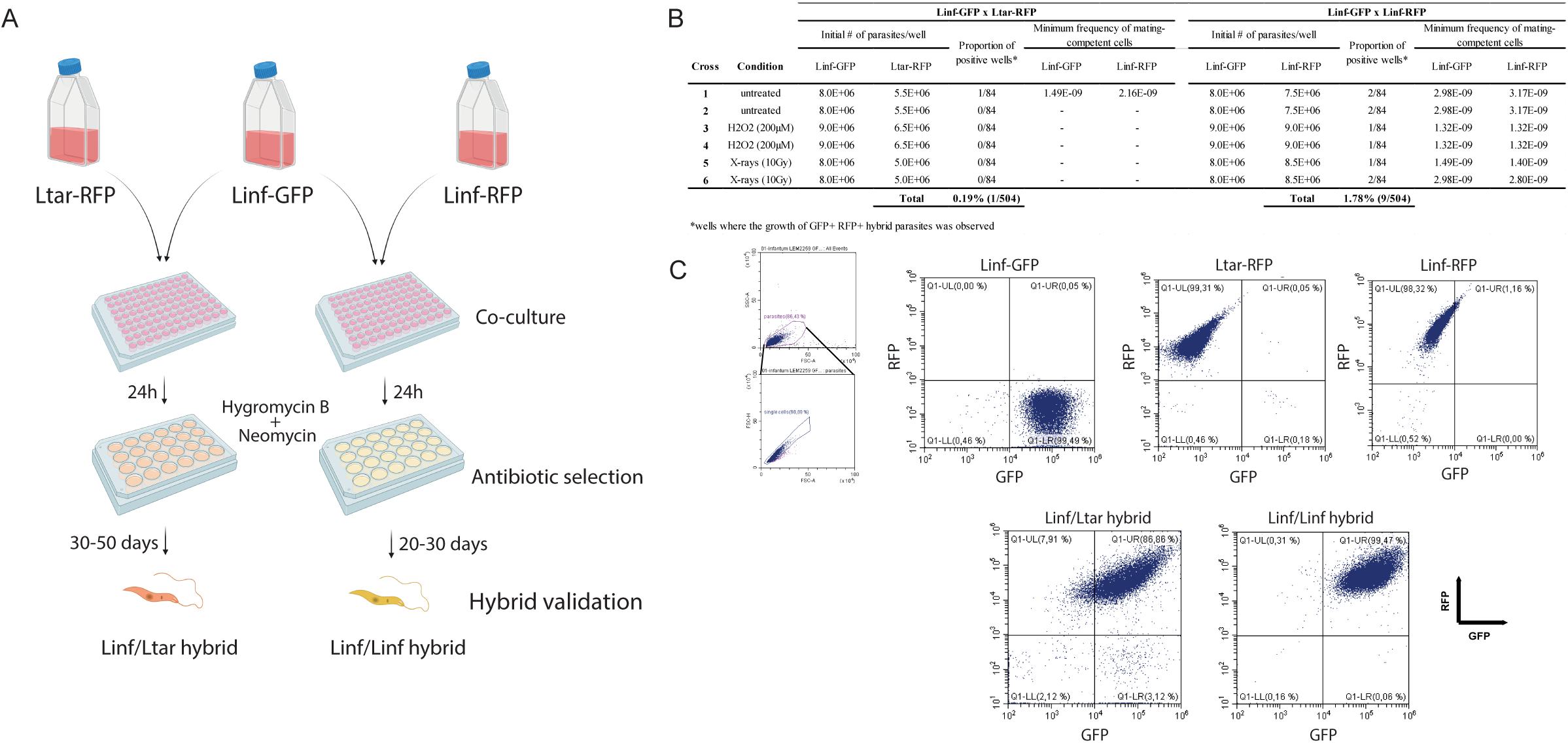
Generation of a Linf/Ltar hybrid by *in vitro* crossing. (**A**) Schematic of the experimental strategy used for the generation of hybrids. (**B**) Summary of the *in vitro* cross series performed (**C**) Validation of the hybrids by flow cytometry on fluorescent markers from the parental lines.

### Assessing uptake and immunostimulatory potential of the Linf/Ltar hybrid using primary mouse and dog macrophages

To analyze the possible immunological impact of the Linf/Ltar hybrid, we next compared the phagocytic uptake of the parental and the Linf/Ltar hybrid strains by primary murine and canine macrophages. In the murine BMDMs, we observed a similar dynamic of infection for the three strains, with a decrease over time observed for the proportion of infected cells (**Figure 2A; Supp. Figure 2**) and the number of parasites/100 cells, with clearance observed for the Linf-GFP parent at 144h PI, and for the Ltar-RFP parent and the Linf/Ltar hybrid at 24h PI (**Figure 2B, Supp. Figure 3**). Of note, a higher parasitic load was observed for the Linf-GFP strain at 4h and 24h PI. For the dog samples, peripheral blood mononuclear cells (PBMCs) were differentiated into macrophages in culture prior to infection with the different parasites. Based on phagocytic tests with zymosan, about 7% of those cells matured into macrophages after 5 days of culture (**Supp. Figure 3**). A higher proportion of dog infected cells was similarly observed with the Linf-GFP strain (from 68.4% at 4h post-infection to 62% at 24h) than with the Ltar-RFP and the hybrid strains (from 62.5% and 57.8% at 4h to 38.4% and 21.9% at 24h, respectively) (**Figure 2C; Supp. Figure 2**) and intracellular parasites were detected at both time points for all strains (**Figure 2D**). Together, these results show Linf/Ltar hybrid and its parental strains are efficiently phagocytosed by both mouse BMDMs and dog PBMCs. However, none sustain infection in mouse BMDMs, though they persist in dog PBMCs at 24h post-infection.

**Figure 2:**
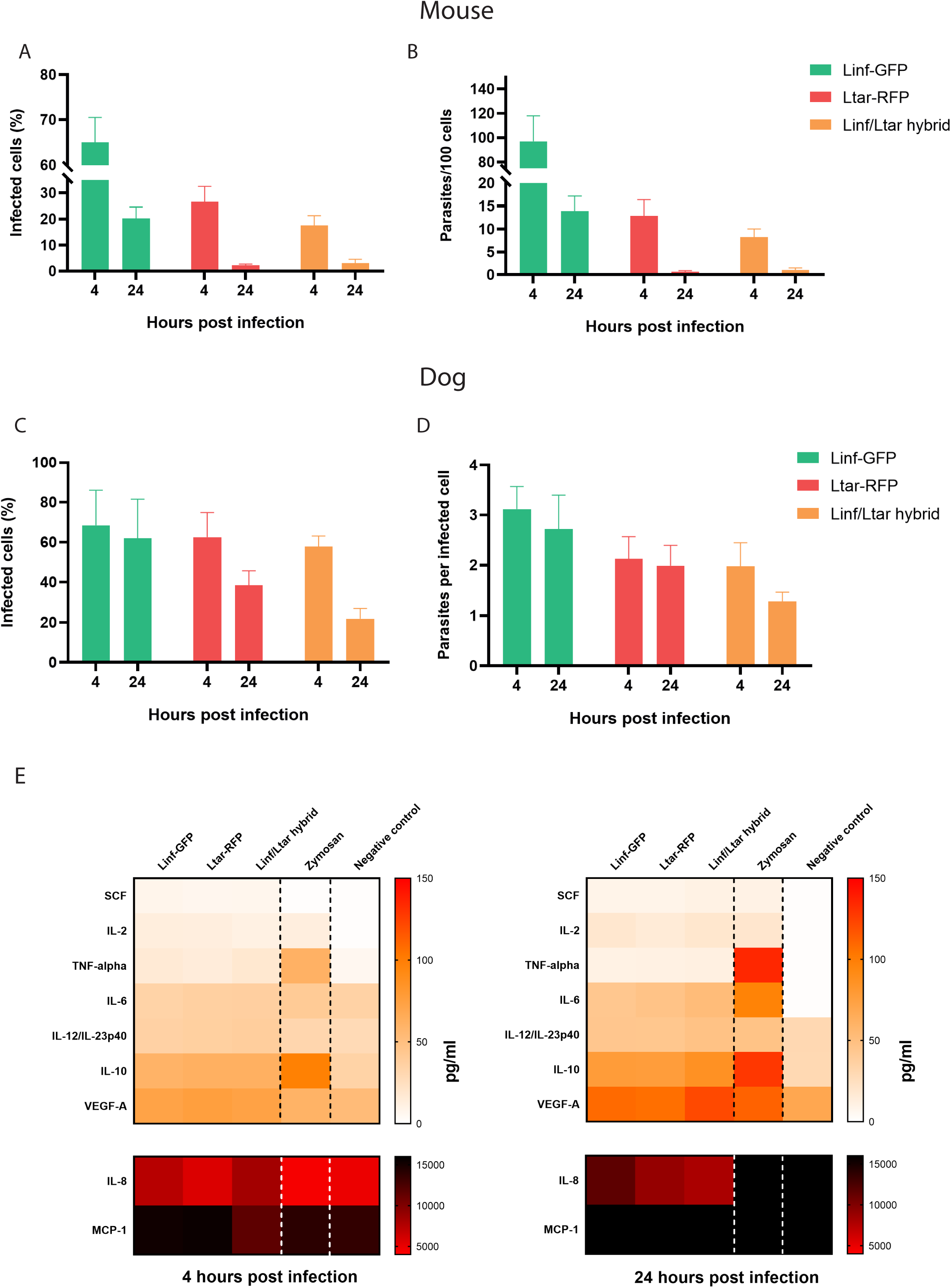
Exposure of mice BMDM and dogs PBMC to the Linf/Ltar hybrid and its parental strains. (**A**) Percentage of infected mouse cells at 4 and 24 hours post-infection. (**B**) Number of parasites per 100 mouse cells at 4 and 24 hours post-infection. (**C**) Percentage of infected dog cells at 4 and 24h. (**D**) Number of parasites per infected dog cell at 4 and 24h. (**E**) Heatmap summarizing the concentrations of each cytokine/chemokine released by the dog cells when infected or not by inf/tar hybrid and its parents, or when exposed to a phagocytic control (zymosan).

We next analyzed cytokine/chemokine production in dog cells infected by parasites or exposed to zymosan for 4h and 24h. Concentrations (pg/mL) of SCF, IL-10, IL-2, VEGF-A, TNF-alpha, IL-8, IL-6, MCP-1 and IL-12 were measured in the culture supernatants (**Figure 2E** and **Supp. Figure 4**). NGF-beta and IFN-gamma were not detectable at any condition. Overall, the infection of dog cells with the Linf/Ltar hybrid or each of its parents led to a similar secretion of SCF, IL-10, IL-2, VEGF, TNF-alpha, IL-8, IL-6 and IL-12 at both time points compared to uninfected cells whereas the concentrations of VEGF and MCP1 stayed unchanged. Compared to *Leishmania* infection, exposure to zymosan induces a higher secretion of IL-10 and TNF-alpha at both time points and of IL-6 at 24h post-infection. The concentration of SCF produced at 4h is on the contrary lower in the cells exposed to zymosan than the concentration detected in the infected cells. These results indicate that infection by the three parasite strains triggers a similar pro-inflammatory response in dog cells, distinct from the response to zymosan beads.

### Comparative genomic analysis of the Linf/Ltar hybrid and its parents

We evaluated the genome ploidy of the Linf/Ltar and Linf/Linf hybrids using Propidium Iodide (PI) staining and flow cytometry, with the diploid parental lines used as normalization controls (**Figure 3A**). All the strains present a typical DNA content pattern: a large peak of cells in the G0/G1 phase and a second, smaller peak of cells in G2/M phase, with a doubled quantity of DNA. The ploidy of the Linf/Ltar hybrid and all the Linf/Linf hybrids analyzed was in each case close to 4n in the G0/G1 phase, suggesting a contribution of each parental genome close to 2n (two complete sets of chromosomes). Whole Genome Sequencing of the single Linf/Ltar hybrid and of seven of the nine Linf/Linf hybrids revealed in each case a biparental inheritance of all the homozygous SNPs that are different between the two parents, resulting in a full-genome heterozygoty (**Figure 3B**), as previously reported for experimental hybrids produced in sand flies and *in vitro* **[15,28–32]**.

**Figure 3:**
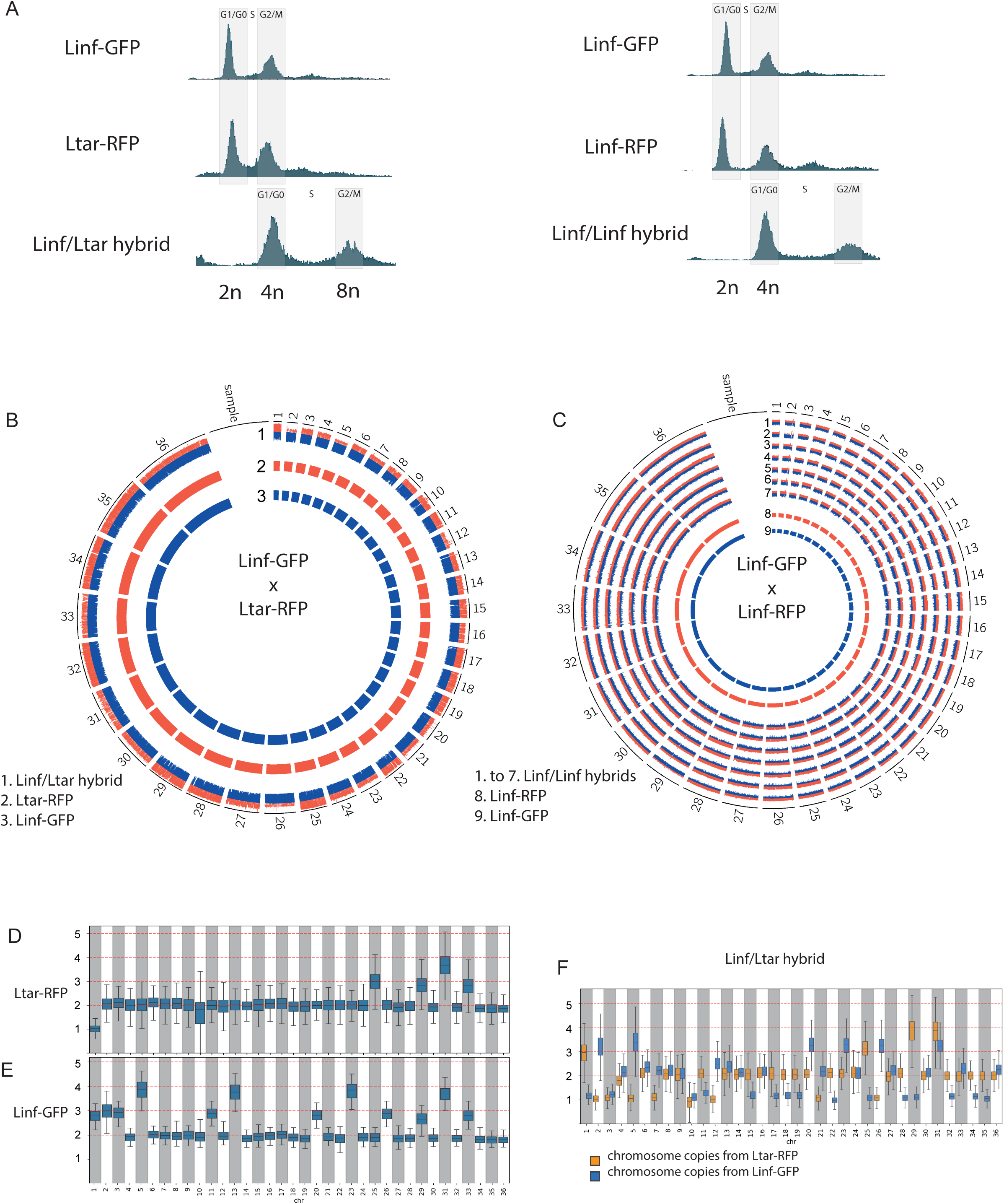
Genomic content analysis of the *in vitro* Linf/Ltar hybrid. (**A**) Observed ploidy of the Linf/Ltar hybrid and one representative Linf/Linf hybrid relative to their parental strains. (**B-C**) Circos plot representation of the inheritance pattern of the homozygous SNP differences existing between their parental strains in the Linf/Ltar hybrid (**B**) and selected Linf/Linf hybrids (**C**). (**D-F**) Individual somies of the Ltar-RFP (**D**), the Linf-GFP (**E**) strains and the Linf/Ltar hybrid (**F).**

We then analyzed the karyotypic profile of the Linf/Ltar hybrid and its parents (**Figure 3D-F**). In both parents most chromosomes were disomic (i.e., with a somy score close to 2) though aneuploidies were observed for chromosomes 1, 10, 25, 29, 31 and 33 for Ltar-RFP (**Figure 3D**), and chromosomes 1, 2, 3, 5, 11, 13, 20, 23, 26, 29, 31 and 33 for Linf-GFP (**Figure 3E**). For most chromosomes, the tetraploid Linf/Ltar hybrid had inherited all the chromosome copies from each of the parents, often resulting in a 2:2 ratio (e.g., chromosomes 6, 8, 9, 13, 14, 16) (**Figure 3F**). However, for several chromosomes, the somy score in the hybrid indicates the loss of one parental copy and thus unequal chromosome contribution. This was observed for the copies of chromosomes 2, 5, 7, 21 and 26 from the Ltar-RFP parent and for the copies of chromosomes 1, 11, 15,17, 18, 19, 22, 25, 28, 32, 24 and 35 from the Linf-GFP parent. Chromosomes 3 and 10 both are disomic in the hybrid, with a single copy of each chromosome coming from each of the parental strains. These results show that the genome of the Linf/Ltar hybrid is not a simple addition of the genetic content of its parents, suggesting that genomic rearrangements during the formation and early growth of hybrids could be important adaptative mechanisms to overcome the genomic shock associated with hybridization **[33,34]**.

### Transcript profiling of the Linf/Ltar hybrid

We questioned whether the Linf/Ltar hybrid transcriptome fully reflects its genomic composition or if certain RNAs are preferentially expressed from one parent. To test this, we isolated and sequenced RNA from axenic promastigotes of Linf-GFP, Ltar-RFP, and the Linf/Ltar hybrid. The RNA reads from Linf-GFP align perfectly to the *L. infantum* reference (98.9% of alignment) but quite poorly to the *L. tarentolae* reference (18.2%). The situation is reversed for the Ltar-RFP strain (18.2% of alignment to the *L. infantum* reference and 98.7% to the *L. tarentolae* reference). 50.3% of the reads from the Linf/Ltar hybrid align to the *L. infantum* reference genome, whereas 66.8% align to the *L. tarentolae* one. When mapping on an in silico-generated hybrid reference (both genomes combined), the alignment rate increases to 99% reads, indicating that both the DNA inherited from the *L. infantum* and from the *L. tarentolae* are transcribed, giving rise to a mixed transcriptome in the hybrid (**Figure 4A**). The PCA plot established from the RNA analysis (mapped on the hybrid reference) also shows a consistent clustering of the experimental replicates from each genotype, with the hybrid clustering in between the Linf-GFP and the Ltar-RFP samples (**Figure 4B**).

**Figure 4:**
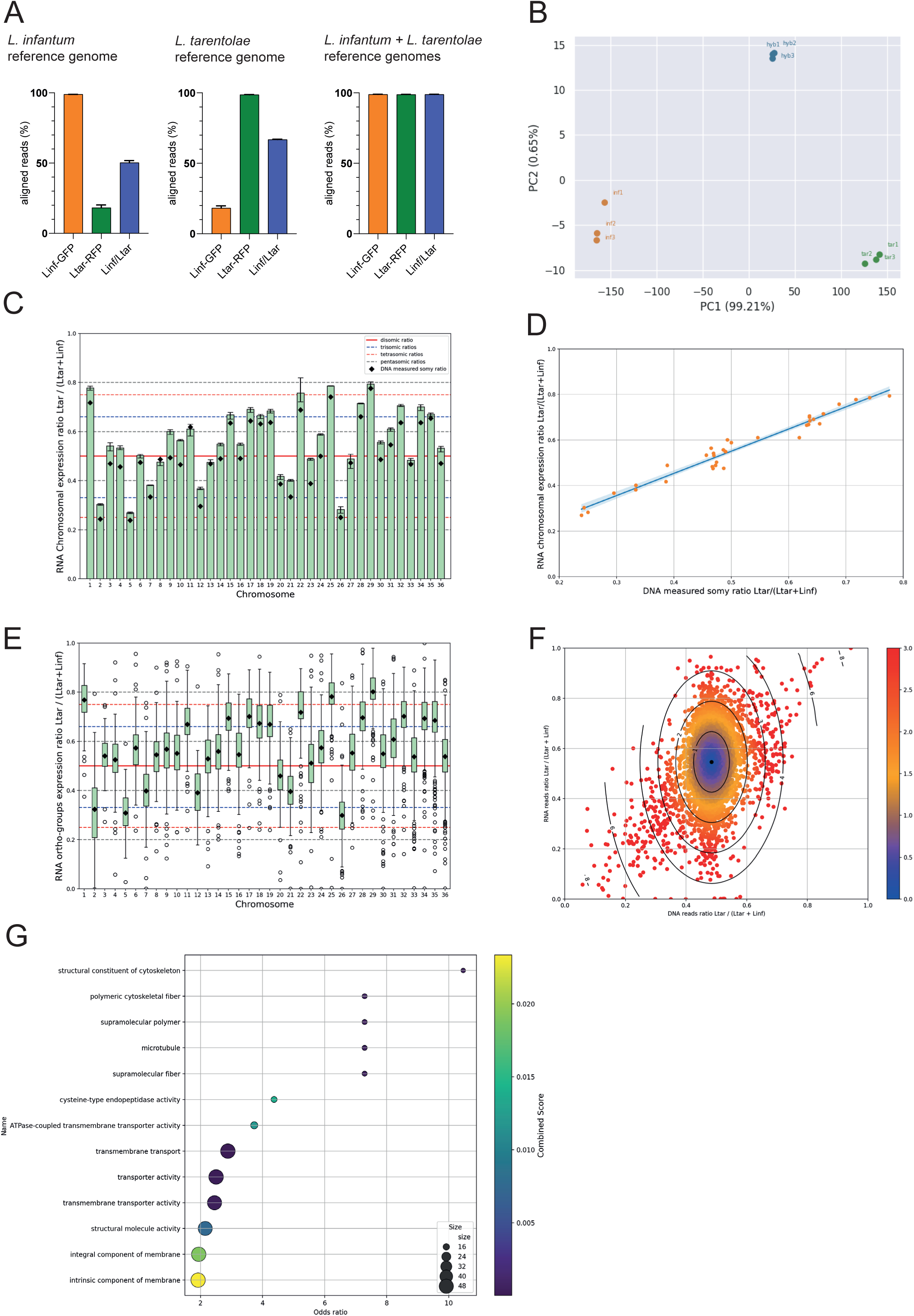
Relative contribution of the *L. infantum* and the *L. tarentolae* genomes to the transcriptome of the Linf/Ltar hybrid. (**A**) Alignments rate of the RNA reads from the Linf/Ltar hybrid and its parents to the *L. infantum* reference genome (left), the *L. tarentolae* reference genome (middle) or both genomes combined (right). (**B**) PCA plots of the 3 replicates of Linf-GFP, Ltar-RFP and Linf/Ltar hybrid transcriptomes, mapped on both genomes combined. (**C**) Proportion of the mRNA reads originating from the *L. tarentolae* genome in the Ltar/Linf hybrid at the chromosome level. The measured somy ratio for each chromosome, calculated with the DNA sequencing data, is indicated by the black diamonds. (**D**) Regression plot between the proportion of reads originating from the *L. tarentolae* genome in the hybrid at the DNA (x-axis) and the RNA (y-axis) levels for each individual chromosome. (**E**) Proportion of the mRNA reads originating from the *L. tarentolae* genome for each orthogroup, when orthogroups are grouped by chromosome. (**F**) Comparison of the proportion of reads originating from the *L. tarentolae* genome for each orthogroup, at the DNA level (x-axis) and the RNA level (y-axis). The z-score for each ortogroup has been color-coded, with z-scores >2 indicating significance. (**G**) Gene Ontology analysis of all orthogroups presenting a z-score >2.

We then investigated the transcriptomic contribution of each parental genome to the overall expression of the hybrid, first at the chromosome level and then at the level of individual genes, by comparing the proportion of reads mapping on the *L. tarentolae* reference (Ltar / (Ltar + Linf)) at the DNA and at the RNA level. At the level of individual chromosomes, we observed a very high correlation between the RNA and the DNA ratios (**Figure 4C-D**), indicating no preferential expression of one parental genome compared to the other. At the gene level, due to differences in annotation between *L. infantum* and *L. tarentolae* genomes that precludes a one-to-one mapping, we first derived orthogroups that allowed us to select 7,347 shared genes for our analysis (see Method section for the selection criteria). Consistent with our chromosome-level analysis, this gene-level analysis showed that most orthologous parental genes are expressed at similar levels (**Figure 4E**). However, from this initial RNA-DNA plot, we observed outliers deviating from the central distribution, indicating that some specific mRNA species could be preferentially enriched from one or the other parental genome.

To better characterize this phenomenon, we plotted the RNA versus the DNA ratios of Ltar/total reads for each orthogroup with a normalization step accounting for the somy score. This adjustment provided a representation of the entire orthogroup dataset, where outliers were more distinctly visible (**Figure 4F**). As explained in the Methods section, we calculated a z-score for each point, which allowed us to identify outliers. The z-scores are color-coded in Figure 4F, with outliers – corresponding to z-scores superior to 2 – represented in red. Different types of outliers can be observed: outliers along the diagonal likely represent gene copy number variations where orthogroups are differently represented between the two genomes, with mRNA abundance reflecting the genomic composition and asymmetric parental gene CNVs. Among those, the top-right corner corresponds to orthogroups where *L. tarentolae* reads are over-represented and over-expressed, while those in the bottom-left corner show the opposite trend. On the contrary, the middle-top quadrant represents orthogroups with balanced *L. tarentolae* / *L. infantum* genomic content but higher expression of *L. tarentolae* mRNA, whereas those in the bottom-middle quadrant present less *L. tarentolae* mRNA for an equal genomic content. Selecting orthogroups with a z-score above 2, we performed GO term enrichment analysis that revealed several enriched GO terms spanning molecular function (MF), cellular component (CC), and biological process (BP) categories. Notably, terms such as “transporter activity,” “structural constituent of cytoskeleton,” and “microtubule-related functions” were identified. A complete listing of these terms is provided in **Figure 4G**. We performed additional GO term enrichment analyses on different categories of outliers, as detailed in **Supplementary Figure 5**.

Overall, our results and analyses indicate that while most transcripts are expressed in a balanced fashion from the parental genomes in the Ltar/Linf hybrid, certain mRNA species are preferentially expressed from one or the other parental genomes. Thus, beyond genomic restructuring, post-transcriptional regulation may serve as an additional mechanism for the establishment of viable hybrid phenotypes that may promote parasite evolvability and fitness gain.

## Discussion

In this study, we demonstrated that *L. infantum* and *L. tarentolae*, two very distinct species belonging to distant clades, can produce sexual hybrid F1 progeny, at least *in vitro*. As far as we are aware, this is the most distant hybrid ever reported, either from field isolates or experimental *in vivo* or *in vitro* production. The occurrence of this hybrid, even as a singular event, indicates that in principle there seems to be no species barrier in *Leishmania* hybridization, even if the frequency of successful hybridization events may depend on the combination of parasite strains and their genetic compatibility. Using a combination of phenotypic, genomic and transcriptomic analyses on a parasite resulting from the hybridization of two highly divergent *Leishmania* strains, our study revealed (i) the potential of the hybrid to stimulate a protective immune response in exposed dogs, (ii) the loss of parental chromosomes as a possible solution to establish genetic compatibility between genetically distinct parents, and (iii) post-transcriptional regulation as an additional mechanism to overcome the genomic shock associated with hybridization. Thus, our study provided valuable insights into the mechanisms involved in the formation of *Leishmania* hybrids and raises important questions on how genetic interference caused by the fusion of two distinct genomes is compensated to establish a viable phenotype.

Our WGS analysis revealed that, despite being close to tetraploid, our Linf/Ltar hybrid does not simply cumulate the genomic content of its diploid parents. We observed the loss of one or several chromosome copies from one or the other parental genomes, which was also detected for other experimental hybrids **[28–32]**. This observation, together with the very low frequency of hybrid formation, clearly emphasizes that hybridization in *Leishmania* is much more than a simple fusion of two cells and addition of their genomes. A previous study based on the analysis of the genomic content of experimental hybrids from the first and second generations demonstrated that hybrid production results from a meiotic-like process **[31]**. However, the identity and more specifically the ploidy of the parental cells fusing is still not established. The hybrids produced in this study were all close to tetraploid, and an important proportion of hybrids previously generated *in vitro* are polyploids (3n or 4n) **[28, 29]**. These ‘young’ hybrid genomes contrast those of field-isolated hybrids that are mainly diploids, which could indicate that fusion of diploid parental cells is the initial step of hybridization, generating a tetraploid hybrid offspring that can later undergo a meiotic reduction of its chromosome content. Another hypothesis would be that diploid hybrids result from a classical meiotic cycle with fusion of haploid gametes (whose existence is so far not demonstrated in *Leishmania*), whereas polyploid hybrids result from the abnormal fusion of a diploid cell with a haploid or a diploid cell. Regardless of the ploidy of the fusing cells, the hybrids resulting from sexual reproduction need to survive, first in the environment where they originated (the sand fly gut) and later in any environment they will encounter. Hybridization, by sexual reproduction or through other types of genetic exchange such as transposable elements or plasmids, leads to a “genomic shock”, a concept first theorized in 1984 by Barbara McClintock **[33, 34]**. This concept originating from plant studies posits that the stress and regulatory interference caused by combining two different genomes can destabilize the genome, potentially leading to significant genetic and epigenetic alterations. This raises the question of the compatibility of parental strains, as the hybridization of genetically more distant organisms could induce a stronger genomic shock, more difficult to overcome. The mechanisms allowing the homeostatic reprogramming of hybrids to establish a viable phenotype are not well understood and probably very different depending on the organisms and the type of genetic exchange. In the case of *Leishmania* products of sexual reproduction, our study indicates that several processes could be involved. At the genomic level, the adjustment of the number of chromosome copies appears as an important mechanism of fitness adaptation, whereas differential mRNA abundance between parental alleles can further buffer possible genetic incompatibilities. *Leishmania* genes are organized in polycistronic units whose transcription initiation is not differentially regulated. The control of gene expression largely occurs at post-transcriptional levels via selective stabilization or degradation of mRNAs **[35]**. Thus, the allelic expression we observed in the hybrid should be controlled by differential mRNA turnover. The molecular mechanisms that can distinguish between different parental mRNA alleles and the underlying structural requirements for differential mRNA turn over eludes us but may involve expression of non-coding RNAs such as snoRNAs, or epitranscriptomic modification of mRNA **[36–38]**. It remains to be determined whether the homeostatic adaptation of hybrids also involves additional regulatory mechanisms, such as translational control, protein turnover, or post-translational modifications.

Finally, a key question in organisms with a facultative sexual reproductive cycle is the role of sex in their evolution. Unlike clonal division, sexual reproduction is a costly and potentially risky process, as it generates entirely new genomes that may be non-viable or less adapted to the host environment than those of the parental organisms. However, this risk can also provide an evolutionary advantage, particularly for organisms facing harsh or fluctuating environments, as the new genome may quickly evolve towards new and more robust adaptive phenotypes, notably in concert with the compensatory mechanisms discussed above. In the case of *Leishmania*, the frequent reports of natural hybrids among field isolates as well as the extraordinary genome plasticity of these parasites indicate that sexual reproduction is more frequent than initially anticipated and may play a central role in both the long-term evolution of *Leishmania* species and their short-term adaptative phenotypes. Further investigation – facilitated by experimental hybrid production - are needed to expand our currently limited understanding of the mechanisms and evolutionary consequences of the *Leishmania* hybridization process.

## Resource availability

Requests for further information and resources should be directed to and will be fulfilled by the lead contact, Isabelle Louradour. The DNA and the RNA data generated in this study have been deposited on the European Nucleotid Archive under the reference E-MTAB-14984 and E-MTAB-14532, respectively.

## Acknowledgements

We gratefully acknowledge Nassim Mahtal and Anne Danckaert from the UTechS Photonic BioImaging (Imagopole), C2RT, Institut Pasteur, supported by the French National Research Agency (France BioImaging, ANR-24- INBS- 0005 FBI (BIOGEN); Investments for the Future), and acknowledge support from Institut Pasteur for the use of the Opera Phenix Plus microscope. We gratefully thank Christophe Ravel from the International *Leishmania* Cryobank and Identification Center in Montpellier, France, for the gift of the *L. infantum* LEM-2259. This work was supported in part by an EMBO Scientific Exchange Grant (V.N.L.F.), by the Agence Nationale pour la Recherche Labex ‘Integrative Biology of Emerging Infectious Diseases’ contract ANR-10-LABX-62-IBEID (S2I program: I.L.), the ANR JCJC program (SFLeisHyb: I.L.), the Roux-Cantarini (I.L.) and the HORIZON-TMA-MSCA-PF-EF post-doctoral programs (SF-Leishyb: I.L.) and the ERC SYNERGY project DecoLeishRN, Grant agreement ID: 101071613 (P.P., G.F.S.). This work was supported in part by the Division of Intramural Research of the NIAID/NIH (T.R.F.).

## Authors contributions

V.N.L.F. designed the project, performed the experiments and analyses and contributed to the manuscript redaction. P.P. performed the macrophages infections. T.C. and T.R.F. analyzed the bioinformatic data and produced the related graphical representations. M.S.L. and J.A.M.R. participated to the elaboration of the project. D.O. and G.S. designed the project and redacted the manuscript. I.L. designed the project, performed the experiments and analyses and redacted the manuscript.

## Declaration of generative AI and AI-assisted technologies

During the preparation of this work, the authors used ChatGPT and Perplexity to improve the readability and ensure correct syntax in the manuscript.

## Supplemental information

**Supp. Figure 1:**
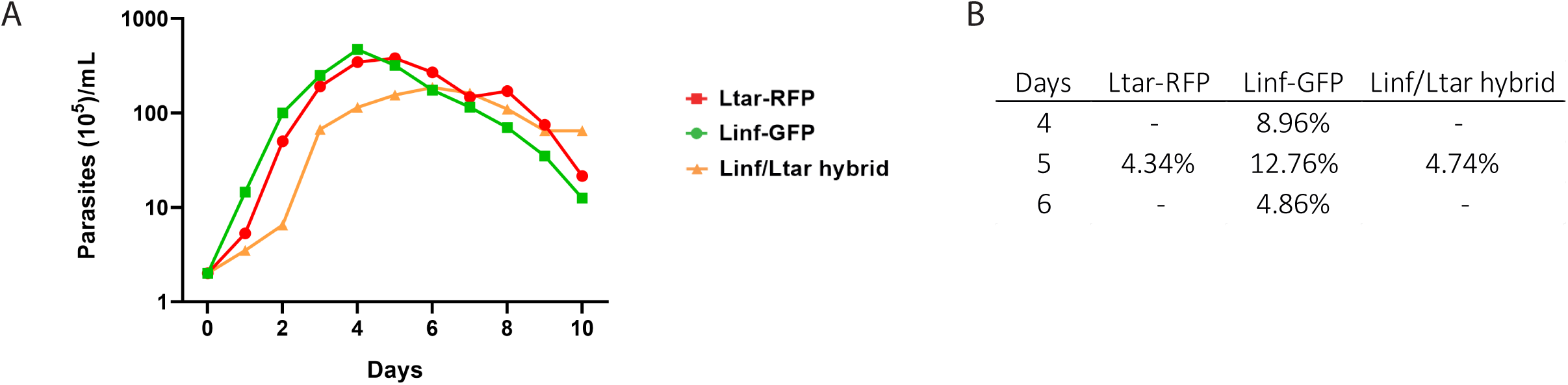
Growth curves (**A**) and metacyclic counts (**B**) of Linf-GFP, Ltar-RFP and the Linf/Ltar hybrid (Related to Figure 2).

**Supp. Figure 2:**
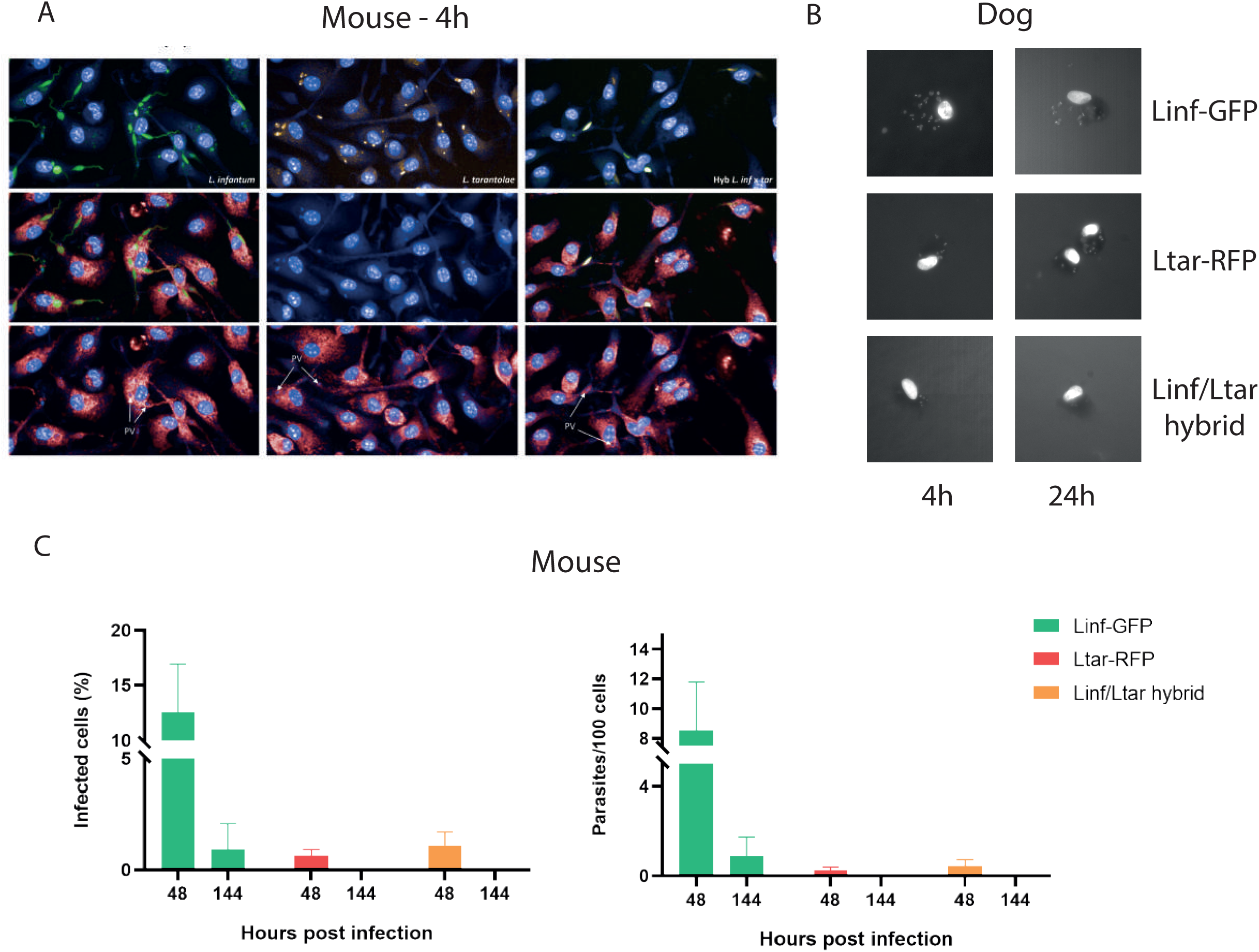
Microscopical pictures of infected mouse BMDM (**A**), infected dog macrophages at 4 and 24h (**B**) and percentage of infected cells and number of parasites per 100 infected mouse BMDMs on late timepoints (i.e. 48 and 144h post-infection) (**C**). (Related to figure 2).

**Supp. Figure 3:**
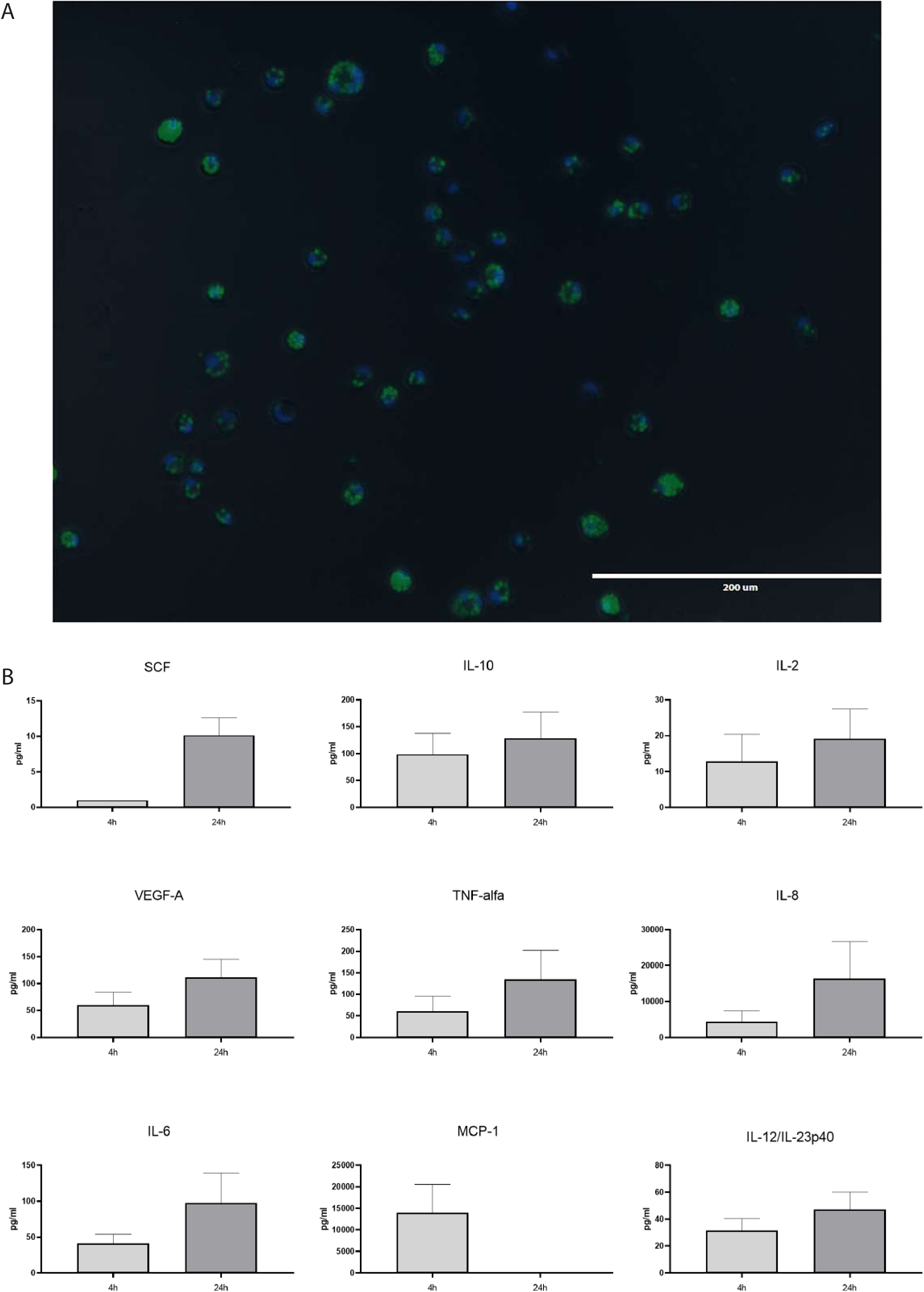
Zymosan data obtained on dog PBMCs (Related to Figure 2). Microscopical pictures (**A**) and cytokine/chemokine concentrations (pg/mL) (**B**) of dog cells incubated with zymosan particles.

**Supp. Figure 4:**
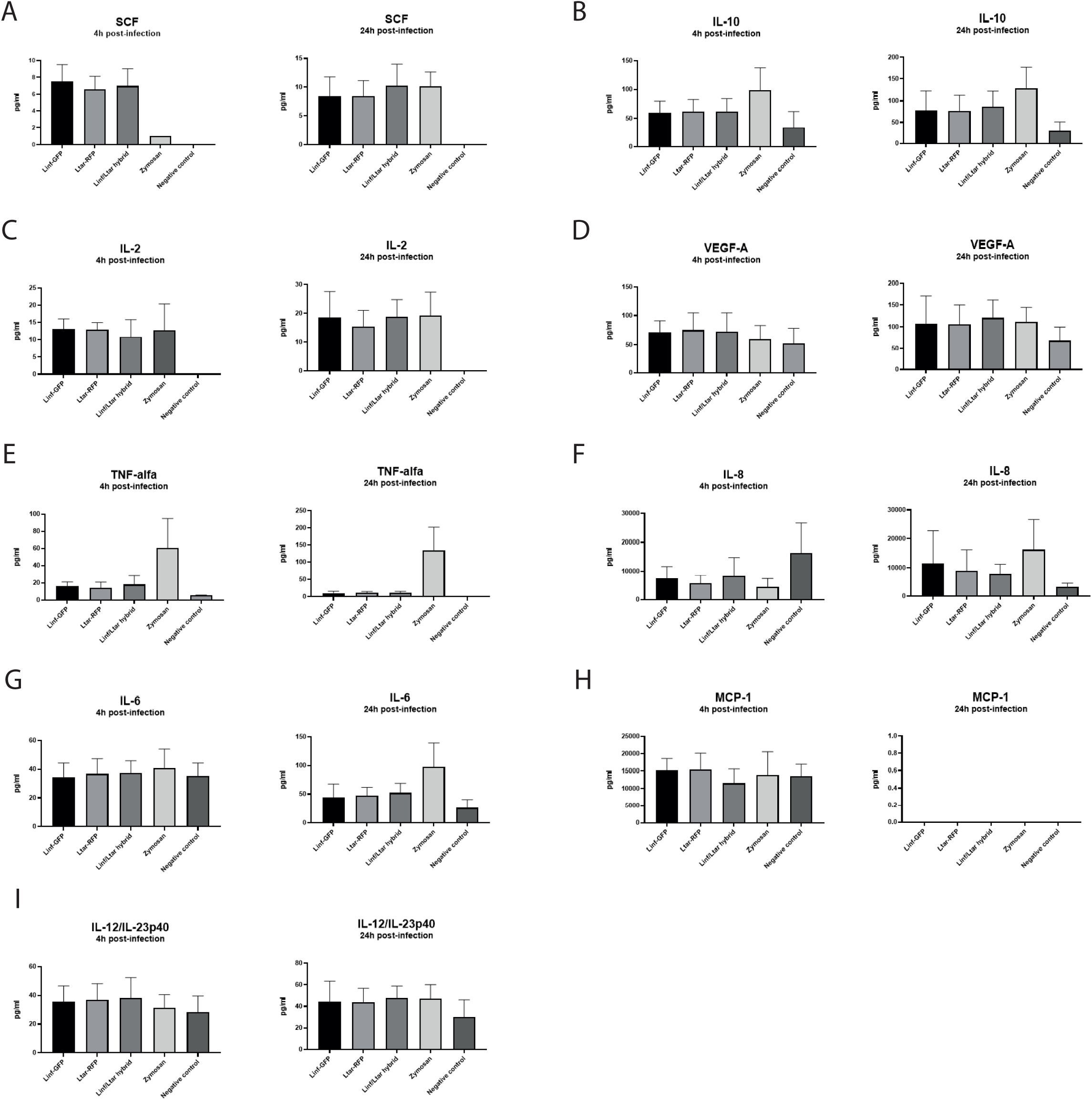
Cytokine and chemokine concentrations (pg/mL) from dog cells after incubation or not with the inf/tar hybrid and its parents or with zymosan particles (Related to Figure 2).

**Supp. Figure 5:**
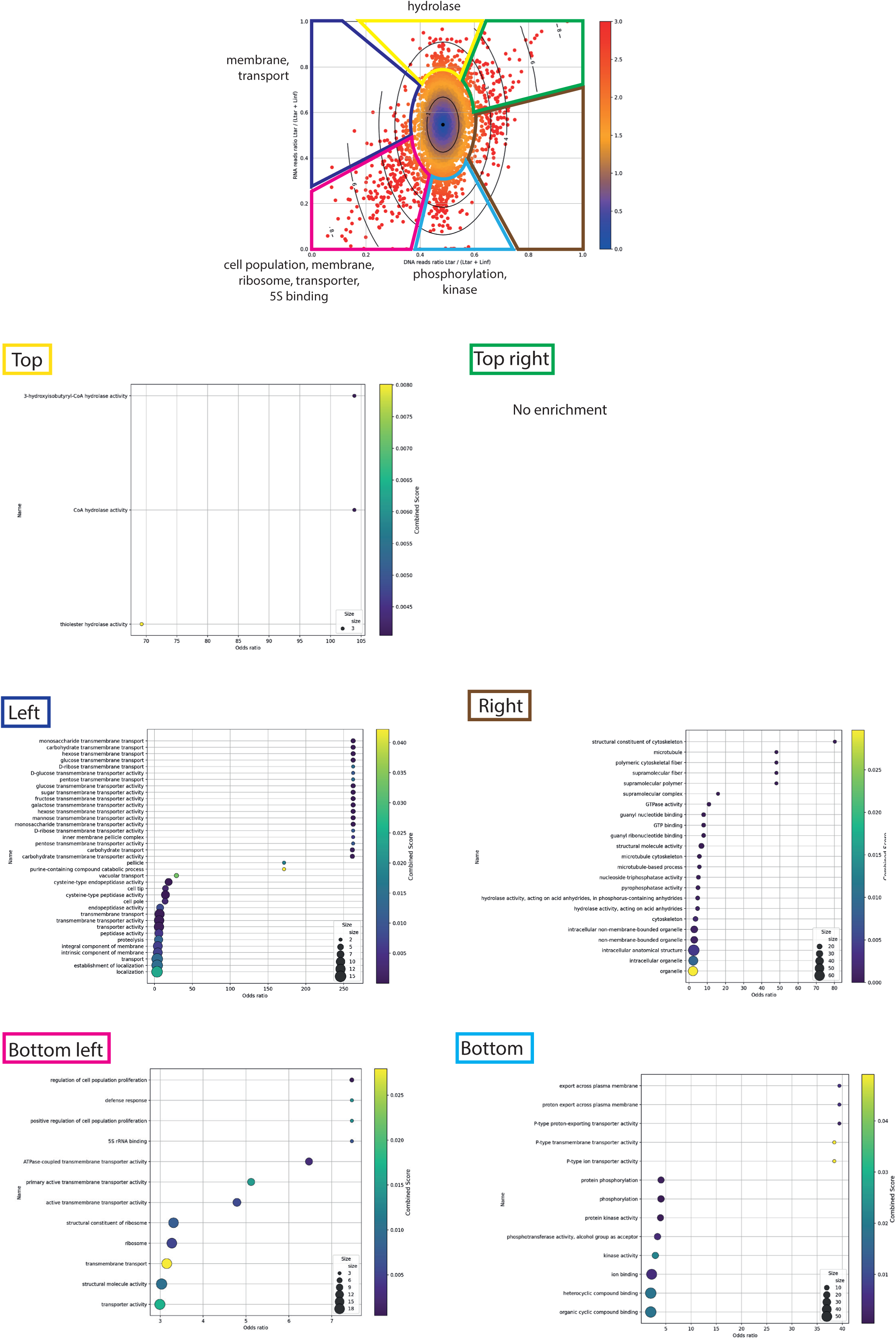
GO term enrichment analyses on different categories of outliers with a z-score>2 (Related to Figure 4).

## Methods

### Parasite strains and *in vitro* hybridization

The following parasite strains were used: *L. infantum* LLM320 (MHOM/ES/92/LLM-320; isoenzyme typed MON-1; **[39]**), *L. infantum* LEM 2259 (MHOM/FR/91/LEM2259, isoenzyme typed MON-1, gift from the International *Leishmania* Cryobank and Identification Center in Montpellier, France) and *L. tarentolae* R011, isolated from a gecko (see **[21]**). The three strains were electroporated using an Amaxa 2D nucleofector (Lonza Bioscience) with integrative expression vectors carrying both a resistance cassette and a fluorescence marker: the pA2-GFP-Neo plasmid, carrying a Neomycin resistance cassette and a GFP fluorescence marker for the *L. infantum* LEM 2259 strain or the pA2-RFP-Hyg vector, carrying a Hygromycin cassette and an RFP reporter resistance for the *L. infantum* LLM320 and the *L. tarentolae* strains **[40]**. The resulting transgenic parasites, referred in the text as Linf-GFP, Linf-RFP and Ltar-RFP, were cloned by limiting dilution and one single clone was used for subsequent use. The promastigotes were cultured at 26°C under classical conditions, in Ld1S promastigote (pro) medium: M199 (ref. 22350, Gibco), 10% FBS, 20 mM HEPES; 100 µM adenine, 2 mM L-glutamine, 10 µg/ml folic acid, 13.7 µM hemin, 4.2 mM NaHCO_3_, 1xRPMI1640 vitamins, 8 µM 6-biopterin, 100 units penicillin and 100 µg/ml streptomycin, pH 7.4. For the *in vitro* generation of hybrids, we used our previously published protocol **[28]**. Briefly, equal volumes of cultures of two parental lines at day 1 post-inoculation were mixed and distributed into 96-well plates at a total volume of 100 µL per well. After 24 hours, each co-culture was transferred to a single well of a 24-well plate containing 900 µL of Ld1S pro medium supplemented with Hygromycin B (25 µg/mL) and Geneticin (Neomycin - 50 µg/mL). Double-drug resistant parasites were passed in new selective medium and analyzed by Flow Cytometry to confirm their expression of GFP and RFP fluorescent markers. Hybrid lines were cloned by limiting dilution, and a single clone from each hybrid line was used for genome and transcriptome sequencing. The daily concentration of promastigotes and the corresponding growth curves were determined by counting a 1/100 culture dilution. The proportion of metacyclic promastigotes was estimated based on the parasites’ morphology and movement (**Supp. Figure 1**).

### Flow cytometry

A CytoFlex Cytometer (Beckman Coulter) was used to observe the parasites fluorescence and to determine their ploidy. The data were analyzed with the CytExpert software V2.4.0.28. The ploidy of hybrid parasites was evaluated using propidium iodide (PI) staining, using the diploid parental lines as normalization controls, as previously described **[28–29]**.

### Ethics statement and animals

The use of mice as source of progenitor cells has been approved by the Institut Pasteur local ethics committee and the animal welfare body (Project DHA-240006). Five-week-old female C57BL/6 were purchased from Janvier labs (CS 4105-Le Genest St Isle 53941 St Berthevin cedex, France) and hosted in A3 animal facility at the Institut Pasteur. All animals were handled under specific, pathogen-free conditions in biohazard level 3 animal facilities (A3) accredited by the French Ministry of Agriculture for performing experiments on live rodents (agreement A75-15-01).

### Mouse BMDM preparation and infection

Murine Bone Marrow-Derived Macrophages (BMDMs) were prepared from the tibias and femurs of three mice as previously described **[41]**. Briefly, bone marrow cells were first plated at a density of 3x10^7^ cells/12 mL in DMEM medium containing 4.5 g/L glucose, 110 mg/L sodium pyruvate, 2 mM L-glutamine, 3.7 g/L NaHCO₃ (Pan Biotech), supplemented with 10% heat-inactivated fetal calf serum (FCS), 50 µg/mL streptomycin, 50 IU/mL penicillin, 50 mM 2-β-mercaptoethanol, 10 mM HEPES and 50 ng/mL of recombinant mouse colony-stimulating factor 1 (rmCSF-1, ImmunoTools) and incubated at 37°C, 7.5% CO2 in tissue culture treated dishes (Falcon 100 mm TC-treated 353003). After overnight incubation, the medium containing the non-adherent cells was collected, diluted in fresh medium containing 50 ng/mL of rmCSF-1 at a concentration of 1x10^6^ cells/mL, and transferred to hydrophobic Petri dishes (Greiner bio-one 664161). After six days of culture, the medium was removed, and adherent cells were incubated with pre-warmed PBS (pH 7.4) containing 25 mM EDTA for 30 minutes at 37°C, 7.5% CO2. BMDMs were detached by gentle flushing, collected, and resuspended in complete medium supplemented with 30 ng/mL of rmCSF-1 at a concentration of 2.5x10^5^ cells/mL. BMDMs were then plated at 2.5 10^4^ cells/well in 96-well plates (PhenoPlate^TM^-96, Revvity) and incubated overnight before infection. The BMDMs were then incubated with 20 parasites/BMDM for each of the different parasite strains or with 3 zymosan particles/BMDM (Invitrogen™ Zymosan A *S. cerevisiae* BioParticles™, Alexa Fluor™ 488 conjugate) as a control for phagocytosis control. The parasites used for infections were prepared by differential centrifugation: promastigote cultures in the late stationary phase (day 5) were centrifuged at 1000 x g for 5 minutes to collect the supernatant containing metacyclic forms. The supernatant was further centrifuged at 3000 x g for 10 minutes, parasites were counted and the culture volume was adjusted to reach a parasite-to-cell ratio of 20:1. Phagocytic and infection assays in murine BMDMs were performed in triplicates with the Linf-GFP, the Ltar-RFP and the Linf/Ltar strains at 4, 24, 48 and 144 hours post-infection. The proportion of infected cells and parasitic load (number of parasites per infected cell) at the different time points were determined using the OPERA Phenix Plus high content imaging system (Revvity) after labeling the cells and parasites. Images from nine fields/well were acquired per replicate and a mean of 2300 cells/well was counted. Image analysis was performed using a dedicated script on the Signals Image Artist (SImA) application. The percentage of infected cells, the number of parasites/infected cells, and the subsequent number of parasites per 100 cells were determined for each time point and each mouse. The results represent the mean values obtained per time point post-infection from the 3 independent mice.

### Immunofluorescence on infected BMDMs

Immunofluorescence was performed on infected murine BMDMs fixed in 4% paraformaldehyde (PFA) in a 96-well plate for at least 30 minutes, followed by a 15 minutes incubation with 50 mM NH₄Cl in PBS. A blocking step was performed with 5% goat serum in PBS-saponin (0.1 mg/mL) for 30 minutes. The cells were then incubated with a rat IgG anti-LAMP 1 (0.5 mg/mL from eBioscience) diluted at 1:100 in PBS-saponin-gelatin (PBS 1x + 0.1 mg/mL saponin + 0.25% gelatin) for 45 minutes, washed three times in PBS-saponin and incubated with a goat anti-rat IgG Fab’2-Alexa 647 (Jackson ImmunoReseach) for 45 minutes. The cells were washed again three times with PBS-saponin, followed by staining with the nuclear dye Hoechst 33342 at a final concentration of 5 µg/mL for 15 minutes. All incubation and washing steps were performed at room temperature. After two final PBS washes, PBS was added to the wells (200 µL/well), and the plates were stored at 4°C until image acquisition.

### Dog PBMC preparation and infection

10 mL of blood/dog collected in heparin-lithium tubes from 3 healthy Beagles were purchased from Marshal BioResources, France. Peripheral Blood Mononuclear Cells (PBMCs) were obtained following the procedure described by **[42]** with modifications. Blood samples were mixed with DMEM medium at a 1:2 ratio (9 mL blood + 18 mL DMEM = 27mL). The mixture was then layered on an equal volume (27 mL) of PANCOLL (1.077 g/mL, Dutscher) and centrifuged at 700 x g for 30 minutes at room temperature to separate the mononuclear cells. The cells of interest, found at the interface of the two layers, were transferred in a new tube, washed twice with 15 mL of warm DMEM medium and centrifuged at 300 x g for 10 minutes at room temperature. The recovered PBMCs were suspended in warm DMEM medium supplemented with fetal bovine serum (FBS), antibiotics (100X Penicillin-Streptomycin) and human recombinant CSF-1 (Proteintech). Cells were seeded at a density of 5x10^5^ cells/mL into 24-well plates for five days and infected on the fifth day with the three parasite lines at a ratio of 10 parasites per cell, or incubated with zymosan particles (3 particles per cell). Zymosan are bio-particles derived from yeast surface cells containing protein-carbohydrate complexes that are recognized by phagocytic cells such as macrophages and are used here as a phagocytic control. The percentage of infected cells and the number of parasites per infected cell were counted daily. All experiments were conducted in technical triplicates for each time point and each dog. At 4h and 24h post-infection, microscopical images were taken after fixation with 4% PFA and nuclei staining with Hoechst 33342.

### ELISA cytokine detection on infected dog PBMCs

The supernatant of the wells containing dog macrophages infected with the parasites or exposed to zymosan was collected and passed onto a 0.22 µm filter at 4h and 24h post-infection and used for quantification of several cytokines and chemokines (IFN-gamma, IL-2, IL-6, IL-8, IL-10, IL-12/IL-23p40, MCP-1, NGF beta, SCF, TNF-alpha and VEGF-A) using the ProcartaPlex™ Canine Cytokine Chemokine Growth Factor Panel 1 11-Plex (Thermo Fisher Scientific, USA) kit, following to the manufacturer guidelines. Briefly, the filtered supernatants from each well were centrifuged at 1400 rpm for 10 minutes at 4°C. A total of 50 μL of standards or samples were added to the 96 well plate containing the capture beads. The plate was sealed and incubated for 2 hours at room temperature with gentle shaking. After incubation and 2 washes, 25 μL of the diluted Detection Antibody Mix was added to each well. The plate was sealed and incubated for 30 minutes with shaking, followed by washing. Next, 50 μL of Streptavidin-PE solution was added to each well, followed by 30 minutes of incubation with shaking. After the final washes, the beads were resuspended with 120 μL of Reading Buffer, and the plate was analyzed using a Bio-Plex instrument. The data were processed to generate a standard curve and calculate cytokine concentrations.

### Reference genomes

The genomic and transcriptomic sequencings were mapped to the *L. infantum* and *L. tarentolae* genomes. For *L. infantum*, both the genome sequence and annotation were obtained from NCBI (accession GCA_900500625.2), consisting of 36 chromosomes and a total size of 32.8Mb. For *L. tarentolae*, the genome sequence and annotation were retrieved from NCBI (accession GCA_009731335.1), consisting of 179 contigs with a total size of 35.4Mb. To study the hybrid samples, we constructed a hybrid reference by concatenating the genome sequences and annotations of *L. tarentolae* and *L. infantum*. While the complete reference genome of *L. tarentolae* was used for mapping, we also constructed a curated version containing only the 57 contigs assigned to known chromosomes. This curated reference accounts for 32.2 Mb, representing 97.3% of the coding genes. The number of coding genes in *L. infantum* is 8,683, while the curated annotation of *L. tarentolae* includes 8,703 coding genes.

### Whole genome sequencing and analysis

DNA was extracted from parasite cultures in log-phase using the DNA blood and tissue kit from MACHEREY-NAGEL, according to the manufacturer’s instructions. Genomic sequencing data were generated using an Illumina NovaSeq 6000 platform, producing paired-end reads with a length of 150 base pairs (2 × 150 bp). The sequencing data exhibits high quality, with an average Phred quality score of 35. Paired-end reads were aligned to the *L. infantum* JPCM5 v.63 reference genome available on TritrypDB (http://tritrypdb.org) using the BWA-MEM aligner v.0.7.17 with default parameters. Single nucleotide polymorphisms (SNPs) were determined using the PAINT software suite designed for studying inheritance patterns in aneuploid genomes **[43]**. PAINT was also used to find and extract the homozygous SNP marker differences between the parental cell lines and to estimate the chromosome copy numbers (somy). Chromosome somies were determined by calculating the normalized median read depth multiplied by 2 (for a diploid genome) using the *ConcatenatedPloidyMatrix* utility with a 5-kb window size. Genomic regions of multiple sequence repeats and high copy number variation (CNV) were filtered out from the analysis by eliminating positions with coverage levels ≥2-fold and ≤0.5-fold the chromosome mean coverage. In the case of the polyploid hybrids (≥3n), the somy values were divided by 2 and multiplied by the ploidy estimated from the DNA content analysis (PI staining) and the parental contribution profile. Allele frequencies of < 0.15, read depth < 10 or represented by < 25% of reads in either forward or reverse direction were filtered out of the analysis. The allelic inheritance of each homozygous parental SNP in the hybrid progenies was determined using the *getParentAllelFrequencies* PAINT utility. The parental allele frequencies were formatted to be compatible with Circos software v.0.69 **[44]**, and inheritance circos plots were generated with 1,257,650 homozygous marker differences between Linf-GFP and Ltar-RFP, and 109,307 homozygous marker differences between Linf-GFP and Linf-RFP cell lines labeled in blue and red, respectively. To estimate the chromosomal somies of the parental strains (figures 3D and 3E), sequencing data from each parent were mapped to their respective reference genomes using the Sequana Mapper pipeline (v1.3.1, https://github.com/sequana/mapper; **[45]**) with default parameters. For the hybrid strain, to assess the contribution of each parent, its genomic data were mapped to the hybrid reference (see reference genome section). Parental contributions were then quantified using the somy-score tool from the Sequana Python library **[45]**. The parental contribution somies were then extracted for subsequent analysis.

### Transcriptomic analysis

Total RNA was extracted using Nucleospin RNA isolation kit (MACHEREY-NAGEL) from the Linf/Ltar hybrid and its parents from three different biological replicates on the second day of growth culture, according to the manufacturer’s instructions. The quality and concentration of RNA were measured by Bioanalyzer DNA1000 Chips (Agilent, # 5067-1504). Transcriptomic sequencing data were generated using an Illumina NextSeq 2000 platform, producing single-end reads with a length of 65bp. The sequencing run yielded 25-80 million reads per sample, with the exception of one hybrid replicate that yielded 8 million reads. The RNA-seq analysis was performed using Sequana v0.17.3 **[45]**. Specifically, we used the RNA-seq pipeline (v0.20.0, https://github.com/sequana/sequana_rnaseq) built on top of Snakemake v6.7.0 **[46]**. Reads were mapped to the aforementioned hybrid reference using Bowtie2 (v2.4.4). FeatureCounts 2.0.0 **[47]** was used to generate the count matrix by assigning reads to features based on the hybrid annotation.

### Orthogroup analysis

To study the relative contribution of each parental genome to the genome and transcriptome of the Lifnf/Ltar hybrid at the gene level, it was necessary to identify the gene orthologs between the two genomes. For that, we used OrthoFinder **[48]** and could assign 16,413 genes (94.4% of the total) into 7,627 orthogroups. We excluded the orthogroups composed of species-specific genes only (e.g. gene duplication on one species, gene loss in the other). Indeed, 94 orthogroups (corresponding to 353 genes) were found to be specific to *L. infantum* and 76 (corresponding to 342 genes) to *L. tarentolae*. We also excluded orthogroups where genes are not located on the same chromosome (within a species or between species), removing 140 additional orthogroups. This left 7,317 orthogroups for the final analysis, encompassing 15,189 genes (87.4% of the total).

### Zscore and Enrichment calculation

FeatureCounts was used to count the number of reads mapped to each gene, applied to both genomic and transcriptomic data. Counts were summed across orthogroups. No normalization was performed since DNA and RNA contributions were calculated as the ratio of *L. tarentolae* reads to the total contribution (*L. infantum* + *L. tarentolae*). The distribution of orthogroups in a two-dimensional RNA-DNA contribution plot revealed a clear relationship centered around equal contributions from both parents. To identify outliers from this distribution, we calculated a z-score based on the Mahalanobis distance, which measures the distance of each point from the meanwhile accounting for the covariance structure of the data. This Mahalanobis-based z-score extends the concept of a standard z-score into two dimensions. Unlike a 1D z-score, which indicates the number of standard deviations a point lies from the mean along a single axis, the Mahalanobis distance considers correlations between dimensions and provides a single, non-negative value for the overall distance from the mean. Despite this difference, there is an approximate relationship between the two: for example, a 1D z-score of 1.5 corresponds roughly to a 2D z-score of 2. This relationship allows us to interpret the Mahalanobis-based z-scores in terms of familiar 1D Gaussian statistics. By using this metric, we identified outliers in the RNA-DNA contribution space for further enrichment analysis. Enrichment analysis of outlier orthogroups was performed using the TriTrypDB database **[49]** with GO enrichment features. Lists of enriched molecular functions, cellular components, and biological processes were retrieved and visualized using custom scripts and Sequana library (e.g. GFF annotation file manipulation). These scripts also included functionality from the Bioservices library to programmatically access the QuickGO website and generate a graph of GO term relationship **[50]**.

